# A SARS-CoV-2 vaccine candidate would likely match all currently circulating strains

**DOI:** 10.1101/2020.04.27.064774

**Authors:** Bethany Dearlove, Eric Lewitus, Hongjun Bai, Yifan Li, Daniel B. Reeves, M. Gordon Joyce, Paul T. Scott, Mihret F. Amare, Sandhya Vasan, Nelson L. Michael, Kayvon Modjarrad, Morgane Rolland

## Abstract

The magnitude of the COVID-19 pandemic underscores the urgency for a safe and effective vaccine. Here we analyzed SARS-CoV-2 sequence diversity across 5,700 sequences sampled since December 2019. The Spike protein, which is the target immunogen of most vaccine candidates, showed 93 sites with shared polymorphisms; only one of these mutations was found in more than 1% of currently circulating sequences. The minimal diversity found among SARS-CoV-2 sequences can be explained by drift and bottleneck events as the virus spread away from its original epicenter in Wuhan, China. Importantly, there is little evidence that the virus has adapted to its human host since December 2019. Our findings suggest that a single vaccine should be efficacious against current global strains.

**One Sentence Summary:** The limited diversification of SARS-CoV-2 reflects drift and bottleneck events rather than adaptation to humans as the virus spread.

## Main Text

Severe Acute Respiratory Syndrome Coronavirus 2 (SARS-CoV-2), the virus that causes COVID-19, is a member of the Coronaviridae family, a diverse group of virus species, seven of which are known to infect humans. Four are considered endemic and typically cause mild upper respiratory illnesses; two of these, NL63 and 229E, are within the alphacoronavirus genus and two, HKU1 and OC43, are betacoronaviruses. The latter genus comprises the three highly pathogenic human coronaviruses, including SARS-CoV-2 as well as Middle Eastern respiratory syndrome (MERS) CoV and severe acute respiratory syndrome (SARS) CoV. SARS-CoV is the most closely related human virus to SARS-CoV-2. While the source, timing and conditions of the zoonotic introduction from SARS-CoV-2 to humans remain unclear, the scale of the pandemic attests to its high transmissibility between humans, with a basic reproduction number R_0_ estimated to be 2.2 (95% CI, 1.4 to 3.9) in Wuhan, China (*1*). In the span of four months, the COVID-19 pandemic has caused an unprecedented global health crisis with significant mortality and socio-economic implications; as of April 23^rd^, 2020, more than two and a half million cases and 184,000 attributable deaths have been reported worldwide (*2-5*) (https://coronavirus.jhu.edu/map.html). SARS-CoV-2 is a single-stranded positive-sense RNA virus, with an approximately 30,000 base pair genome. The genome is split into ten open reading frames (ORF) that include fifteen non-structural proteins and four structural proteins. The latter category includes the Spike (S), Membrane (M), Envelope (E) and nucleocapsid (N). The S protein is the basis for most candidate vaccines, as it mediates virus attachment and entry to host cells and is the target of neutralizing antibody responses (*6-8*). The S protein is cleaved into two subunits, S1 and S2: the former contains the receptor binding domain (RBD), which enables the virus to attach to the angiotensin-converting enzyme 2 (ACE2) receptor on host cells.

Developing a vaccine against SARS-CoV-2 is a high priority for preventing and mitigating future waves of the pandemic (*9*). Vaccine candidates typically include an insert that corresponds to one or more virus antigens, either derived computationally or from one or multiple strain(s) sampled from infected individuals. The first viral sequence derived during the COVID-19 outbreak, Wuhan-Hu-1 (GISAID accession EPI_ISL_402125), was published on January 9, 2020. As many vaccine programs initiated at that time, it is likely that this SARS-CoV-2 sequence, sampled in December 2019 in Wuhan, China, will be the foundation for a number of vaccine candidates. Compared to other RNA viruses, coronaviruses have a more complex molecular machinery resulting in higher replication fidelity. The mutation rate of SARS-CoV was estimated at 9.0×10^−7^ substitutions per nucleotide per replication cycle (*10*), a rate that is much lower than that of most RNA viruses (1×10^−3^ to 1×10^−5^ substitutions per nucleotide per replication cycle) (*11*), indicating the limited propensity of coronaviruses to diversify. Nonetheless, as the virus spreads more widely, it is important to monitor the introduction of any mutations that may compromise the potential efficacy of a vaccine candidate based on the first sequence.

We can obtain insights about the transmission pathways and temporal spread of a virus by analyzing viral sequences sampled from individuals who became infected. Changes in allele frequencies across sequences indicate the fitness of the virus over time, as it adapts to its host. Increasing fitness will result in pervasive mutations at specific sites that improve virus transmissibility, whereas mutations in a neutral evolution context will be minimal and stochastic. Indicators of viral evolution have been shown to be robust predictors of transmission dynamics for several pathogens, such as Influenza (*12*), Lassa virus (*13*), and Ebola (*14*). Typically, the evolution of a virus is driven by genotypic and phenotypic changes in its surface protein. In the case of SARS-CoV-2, mutations in the protein S, which mediates cell entry, are most likely to confer fitness to the virus as it adapts to humans. However, adaptive changes can occur in structural and non-structural proteins, and these changes, as well as different patterns across structural and non-structural proteins, may provide insights into the near- and long-term evolutionary dynamics of SARS-CoV-2, as it spreads in humans.

To characterize SARS-CoV-2 diversification since the beginning of the epidemic, we aligned 5,724 independent SARS-CoV-2 genome sequences (average length of 29,874 nucleotides) isolated from 68 countries (**Figure 1A**) (*15*). There were 4,678 polymorphic sites. Only 1,231 (4.19%) showed mutations in two or more sequences (**Figure 1B**). The mean pairwise diversity across genomes was 0.08%, ranging between 0.01% for ORF7b to 0.11% for the nucleocapsid (N). Hamming distances across all genomes showed a narrow distribution with a median of nine nucleotide mutations (**Figure S1**). A phylogenetic tree reconstructed based on all genome sequences (*16*) reflected the global spread of the virus: samples from the first six weeks of the outbreak were collected predominantly from China and were more basal in the tree (**Figure 1C**). As the epidemic has progressed, samples have been increasingly obtained across Europe and from the United States (**Figure 1A, 1C**). The tree shows multiple introductions of different variants across the globe with introductions from distant locations seeding local epidemics, where infections sometimes went unrecognized for several weeks and allowed wider spread (*17*). The tree topology shows minimal structure, even at the genome level, indicating that SARS-CoV-2 viruses have not diverged significantly since the beginning of the pandemic.

**Figure 1.**
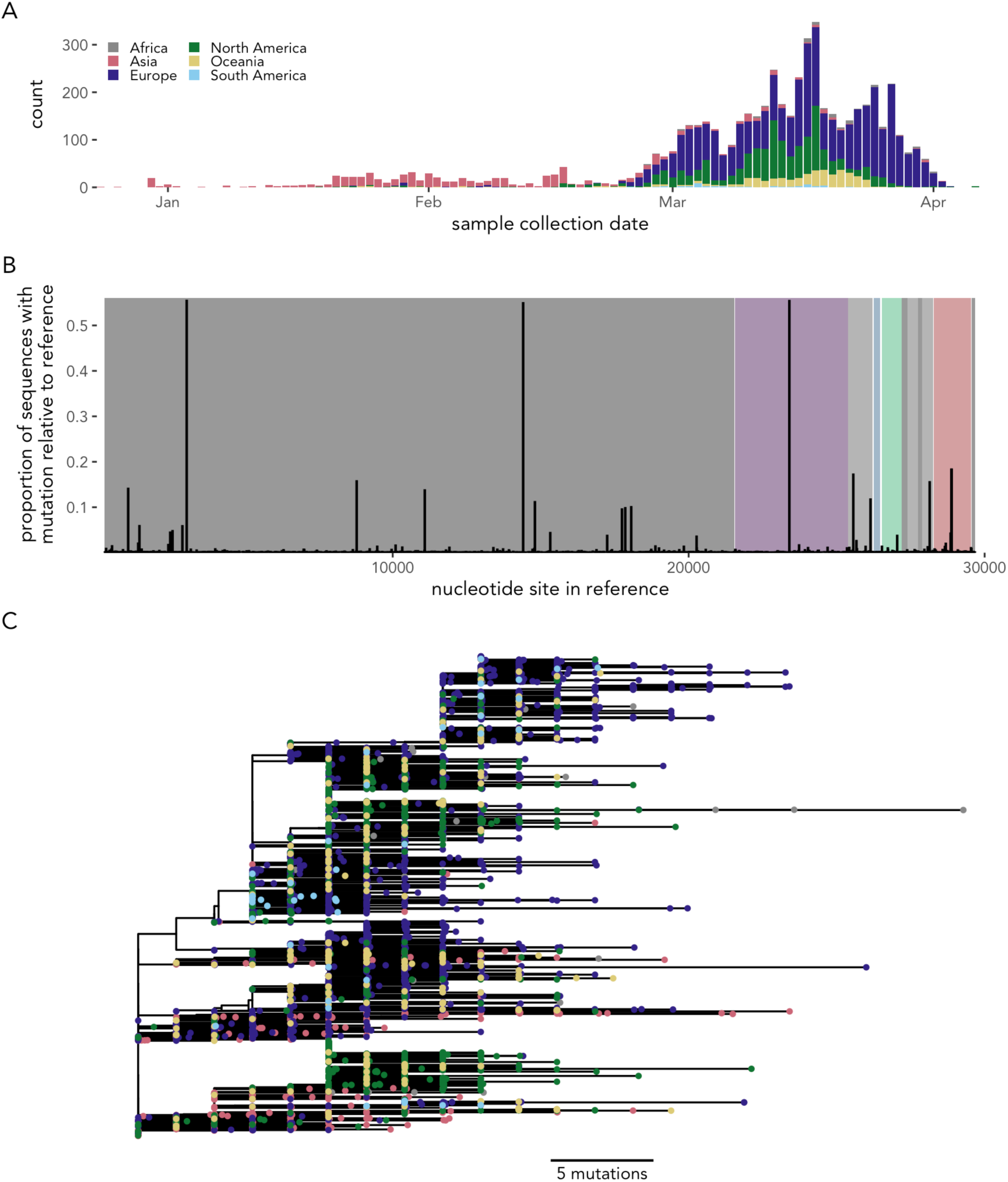
SARS-CoV-2 diversity across 5,724 genomes. (A) Distribution representing the location and date of sample collection. (B) Location and frequency of mutations across the genome. Proportion of sequences which have a mutation at the nucleotide level compared to the reference sequence, Wuhan-Hu-1 (EPI_ISL_402215). Open reading frames are shown in gray for non-structural proteins and in color for structural proteins (Spike, S: purple; Envelope, E: blue; Membrane, M: green; Nucleocapsid, N: red). (C) Global phylogeny of 5,579 independent sequences. Tree reconstructed from all near-full length sequences (tips more divergent than expected from their sampling date were removed). The tree was rooted at the reference sequence, Wuhan-Hu-1, and tips colored by collection location. The scale indicates the distance corresponding to five mutations.

Even with minimal divergence, we verified whether specific sites in the genome were selected to diversify. We used likelihood-based, phylogenetically informed models assuming branch-specific substitution rates (*18*) and identified eight sites under diversifying selection (i.e. with a non-synonymous/synonymous substitution rates ratio dN/dS > 1), including four sites in S: sites 5, 50, 614 and 943 (**Figure 2A**). Three of these four sites rarely varied (with the variant amino acid found in <1% of sequences) and only transiently diversified thus far during the epidemic. In contrast, site 614 changed rapidly: the D614G mutation was introduced into Europe in January and became dominant on that continent (**Figure 2B**) and then globally within two months (found in 55% of sequences) (**Figure S2**). Overall, at least two thirds of sites across the genome were under purifying or neutral selection (dN/dS<1): S=85.9%, E=68.0%, M=74.8%, N=65.6% (**Figure 2C**) (*19*). While dN/dS ratios varied across genes, ranging from 0.32 (M) to 1.19 (ORF6), they were not significantly different between structural and non-structural genes (Mann-Whitney test, P=0.32) (**Figure 2C**). Per-lineage mutation rates per site, calculated on internal nodes, also showed no differences in mean values (P=0.90) or distributions (K-S test, P=0.89) between structural and non-structural genes (**Figure 2D**). Despite notable differences across structural genes, all fell in the range of other RNA viruses (**Figure 2E**) (*20*). Mutations were disproportionately neutral: all branches evolved under neutrality for >65% of sites; and half of all branches evolved under neutrality for >83% of sites (**Figure 2F**) (*18*).

**Figure 2.**
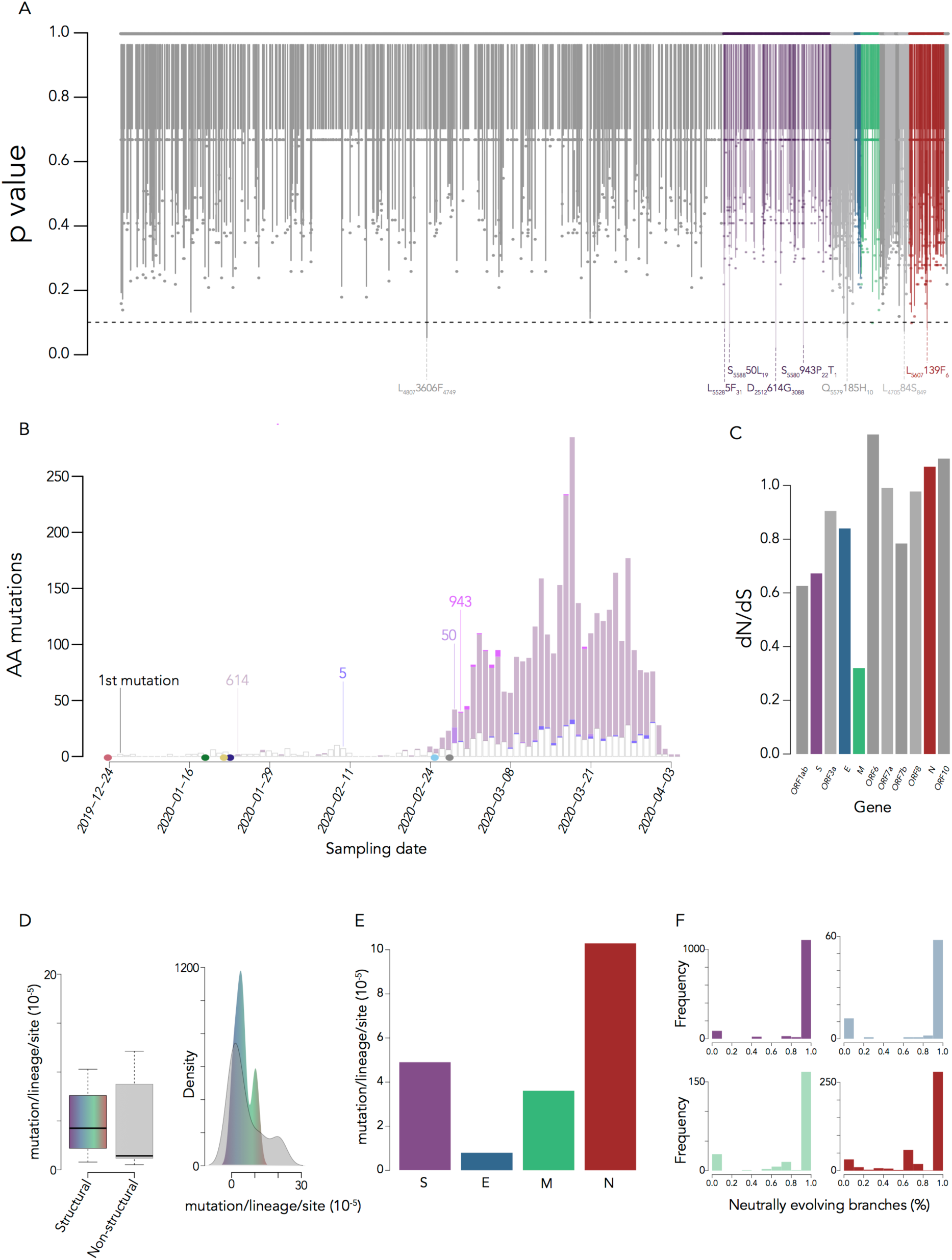
Evolution across the SARS-CoV-2 genome. (A) Barcode plot of p-values for each site across the SARS-CoV-2 genome estimated from likelihood-based models of selection. Sites with P<0.1 (dashed line), indicating evidence of diversifying selection, are shown with the proportion of variant sites. (B) Number of mutations in the Spike protein across all sequences sampled at each time point. The date of first sampling for each continent (filled circles) and of first appearance of specific variants (lines) are shown. (C) Global estimates of non-synonymous/synonymous rates for each gene inferred from a reversible MG94 codon model. (D) Boxplot and density plot of mutation rates for structural and non-structural genes. (E) Mutation rates for structural genes. (F) Frequency of branches evolving under neutrality across sites within each gene.

While there was only limited evidence of diversification at selected sites, we also assessed whether the globally circulating viral population differentiated from the initial population sampled in Wuhan. To do so, we used two measures of population differentiation, the G_ST_ and D statistics, which characterize changes in allele frequency across populations and can show fitness differences within subpopulations (*21-23*). We compared 353 genomes sampled from the initial outbreak in Wuhan, China with 5,280 genomes sampled subsequently across the globe. Genetic distances between two subpopulations can range between 0 and 1, indicating no and complete differentiation, respectively. Although distances varied across genes, the genetic distance between these subpopulations was small, indicating little differentiation between the initial outbreak and its global spread (**Figure 3A**). (Elevated estimates can also be artifacts associated with high mutation rates (**Figure 2E**) (*24*)).

**Figure 3.**
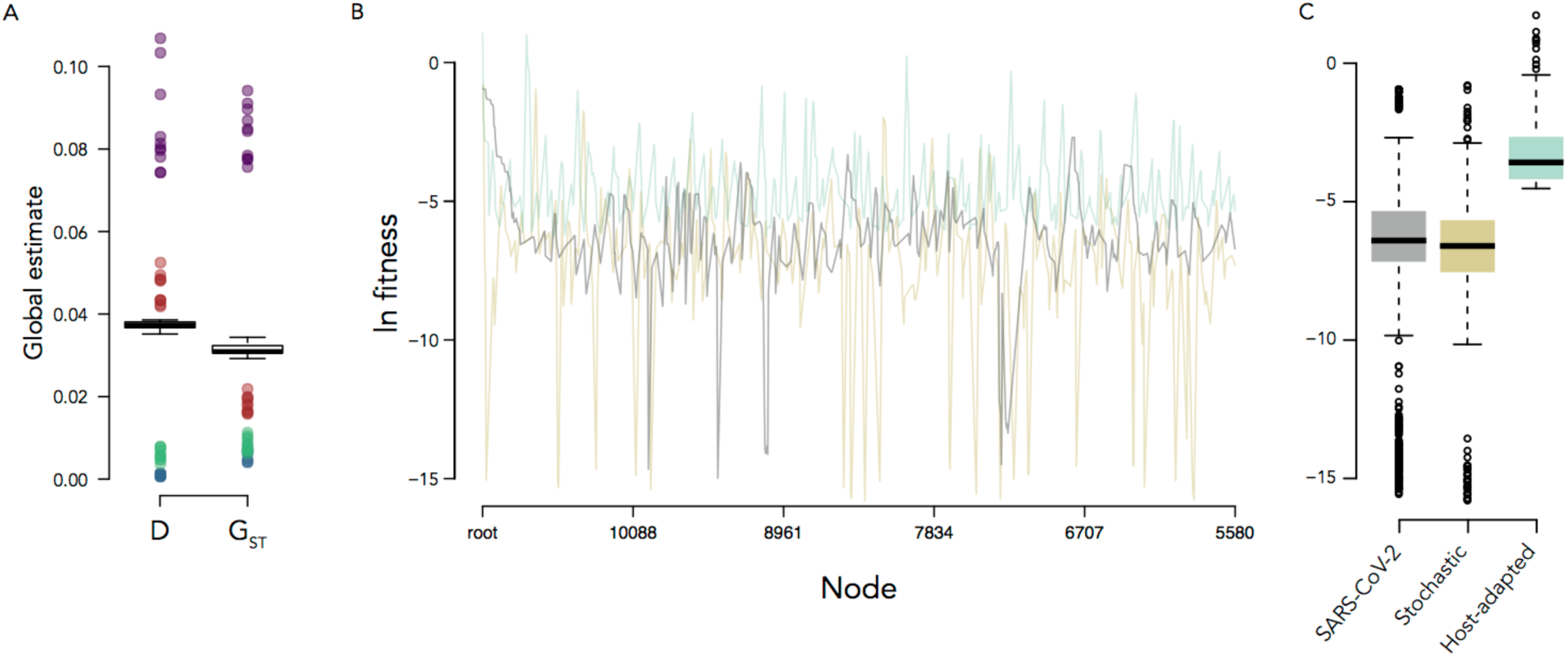
No evidence of host adaptation among SARS-CoV-2 sequences. (A) Bootstrapped global estimates of Jost’s D and Nei’s G_ST_ across all genes and estimates from bootstrapped samples for each gene separately (Spike, S: purple; Envelope, E: blue; Membrane, M: green; Nucleocapsid, N: red). (B) Mutational fitness for subtrees at each internal node for the SARS-CoV-2 S gene tree (grey), neutral fitness model (gold), and increasing fitness model (teal). (C) Average mutational fitness calculated over down-sampled phylogenies.

Signatures of host adaptation can also be seen in the branching patterns of viral phylogenies. Bursts in transmissibility are emblematic of increases in relative viral fitness and are reflected in imbalances in the phylogeny, which can be estimated at each internal node (**Figure S3**) (*25-27*). We estimated mutational fitness (*28, 29*) at each internal node of the SARS-CoV-2 phylogeny reconstructed from the *spike* gene and compared the distribution of estimates through time to phylogenies simulated under models of neutral and increasing fitness (*i.e.*, host-adapted). The distribution adhered to expectations of the neutral model and deviated significantly (Mann-Whitney test, P<1e-3) from those of increasing fitness (**Figure 3B**). Similar results were obtained when the SARS-CoV-2 and simulated phylogenies were iteratively down-sampled by 50% (**Figure 3C**). Therefore, we concluded that the sample size of sequences was sufficient to test for adaptation. Together, the SARS-CoV-2 population and phylogeny showed no evidence that the global spread of SARS-CoV-2 was related to viral fitness effects.

Last, we evaluated two typical vaccine design strategies, which rely either on sampled sequences or on computationally derived sequences that cover the diversity seen across circulating sequences and are in theory optimal compared to an individual isolate (*30*). We inferred the most recent common ancestors (MRCA) of SARS-CoV-2 S sequences sampled from Wuhan within the first month of the epidemic, from all currently circulating SARS-CoV-2 sequences, and from those corresponding to closely related sequences sampled from pangolins (n=9) and a bat. There were 17 mutations between the human MRCA and the human-bat MRCA and 41 mutations between the human MRCA and human-pangolin MRCA. Overall, three segments in S reflected significant variability across species (AA 439-445, 482-501, 676-690) (**Figure 4A-B, Figure S4**) (*31*). In contrast, when considering only human sequences, SARS-CoV-2 diversity was limited: both MRCAs (derived from early Wuhan sequences or from all circulating sequences) were identical to the initial reference sequence Wuhan-Hu-1 (**Figure 4C**). Comparing these sequences to the consensus sequence derived from all of the sequences sampled to date, there was only one mutation: D614G (**Figure 4C**). **Figure 4D** illustrates that mutations found across circulating sequences are extremely rare: beside D614G (found in 55% of sequences), the next most frequent mutation is found in 0.87% of sequences, with sequences sampled from infected individuals on average 0.69 mutations away from the consensus sequence (consisting of 0.11 synonymous and 0.57 non-synonymous mutations). Across the genome, there were on average 11.46 nucleotide mutations per individual genome when compared to the consensus, with only two other mutations aside from D614G found in >50% of sequences. Focusing on the RBD, we found no evidence that mutations could affect binding to the ACE2 receptor, as only two shared mutations were identified at contact sites with the ACE2 receptor: a non-synonymous mutation (G476S) found in 8 sequences (0.15% of sequences) and a synonymous mutation found in 2 sequences at position 489 (0.04% of sequences).

**Figure 4.**
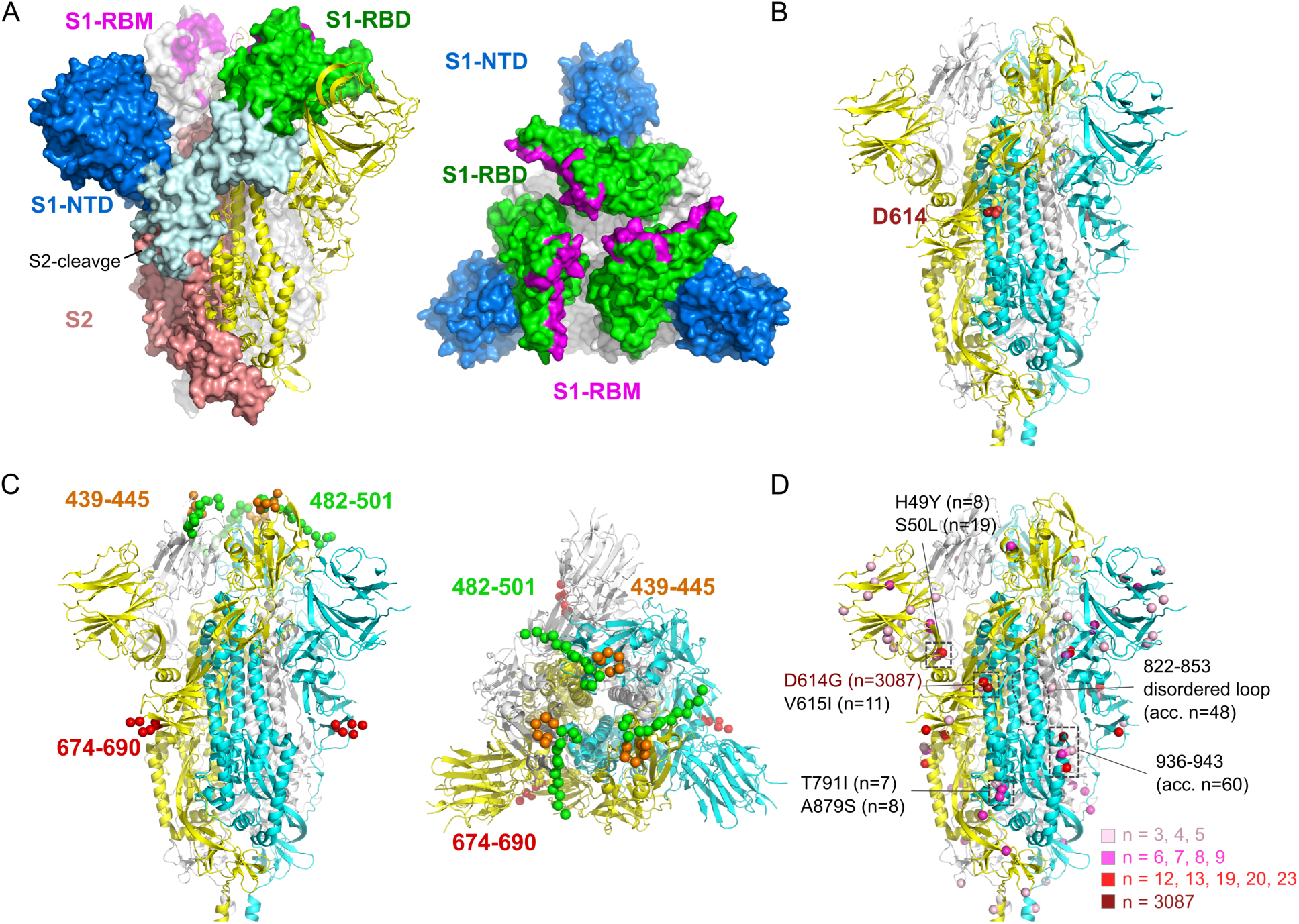
Mutations across SARS-CoV-2 Spike sequences. (A) Structure of SARS-CoV (5×58) (shown instead of SARS-CoV-2 for completeness of the RBM). (B-D) The three protomers in the closed SARS-CoV-2 Spike glycoprotein (PDB code 6VXX) are colored in yellow, cyan and white. Sites with mutations are shown as spheres. (B) Near-identity of potential vaccine candidates, the MRCA and Wuhan-Hu-1 reference sequences were identical while the consensus derived from all circulating sequences showed a mutation (D614G). Site 614 is located at the interface between two subunits. (C) Sequence segments that differed between human and pangolin or bat hosts. Amino acid segments 439-455 and 482-501 impact receptor binding while the 574-690 segment corresponds to the S2-cleavage site. (D) Sites with shared mutations across 5,607 SARS-CoV-2 circulating sequences. The colors of the spheres correspond to the number of SARS-CoV-2 sequences that differed from the Wuhan-Hu-1 sequence (GISAID: EPI_ISL_402125, GenBank: NC_045512). Mutations that were found only in one or two sequences were not represented.

There remains an urgent need for a SARS-CoV-2 vaccine as a primary countermeasure to contain and mitigate the spread of COVID-19. The virus’s surface S protein makes an attractive vaccine target because it plays a key role in mediating virus entry and is known to be immunogenic (*32*). Neutralizing antibody responses against S have been identified in SARS-CoV-2 infected individuals (*8*) and several clinical trials for a SARS-CoV-2 vaccine will test S as an immunogen. While we focused on the S protein, our comparative analyses of other proteins yielded similar conclusions: a randomly selected SARS-CoV-2 sequence could be used as a vaccine candidate given the typical similarity of any sequence to the computationally derived optimum sequence (as defined by the MRCAs or consensus sequence based on all circulating strains). Vaccines developed using any of these sequences should theoretically be effective against all circulating viruses.

Our analyses showed limited evidence of diversifying selection, with comparable mutation and evolutionary rates in structural proteins versus non-structural proteins (under a selection paradigm, structural proteins which are essential for viral entry and the target of the host immune response would have higher rates than the non-essential proteins), low estimates of genetic differentiation following the initial outbreak, and phylogenetic patterns adhering to a neutral process of evolution. These results suggest that SARS-CoV-2 has not, as yet, adapted to its host as it is spreading globally. Rather, our findings illustrate that epidemiological factors primarily mediated the global evolution of SARS-CoV-2. While certain mutations have increased in frequency, these were likely exported to under-infected areas early in the outbreak and therefore given the opportunity to spread widely (founder effect). For example, in January 2020, a virus carrying the D614G mutation, which had not then been observed among sequences from China, was transmitted to Germany. Host and environmental factors permitted the establishment of a sustained cluster of infections that propagated this mutation until it became dominant among European sequences and then globally (**Figure S2**). There is no evidence that the frequent identification of this mutation is caused by convergent selection events that would have occurred in multiple individuals. The D614G mutation is at the interface between two subunits and thereby would not be expected to be part of a critical epitope for vaccine mediated protection (**Figure 4**). However, an S612L mutation, in the same region as the D614G mutation, has been observed after passaging a MERS-CoV isolate in the presence of two antibodies (in 5/15 clones after 20 passages) (*33*). The rare presence of the S612L mutation among MERS-CoV sequences (2 of 473 sequences from humans) indicates that it is unlikely to be a common escape pathway. Nonetheless, its occurrence *in vitro* warrants further work to evaluate whether the close-by D614G mutation in SARS-CoV-2 impacts protein stability, conformational changes, infectivity or recognition of a distal epitope. While the possibility of a D614G escape pathway is unknown, mutations in the RBD constitute a likely path to escape antibody recognition, as described for Influenza (*34, 35*). Importantly, we found no mutation in the RBD that was present in more than 1% of SARS-CoV-2 sequences; such rare variants are unlikely to interfere with vaccine efficacy. These results indicate that so far SARS-CoV-2 has evolved through a non-deterministic, noisy process and that random genetic drift has played the dominant role in disseminating unique mutations throughout the world. Yet, it is important to note that the virus was only recently identified in the human population – a short time frame relative to adaptive processes that can take years to occur. Although we cannot predict whether adaptive selection will be seen in SARS-CoV-2 in the future, the key finding is that SARS-CoV-2 viruses that are currently circulating constitute a homogeneous viral population. Viral diversity has challenged vaccine development efforts for other viruses such as HIV-1, Influenza or Dengue but these viruses each constitute a more diverse population than SARS-CoV-2 viruses (**Figure S5**). We can therefore be cautiously optimistic that viral diversity should not be an obstacle for the development of a broadly protective SARS-CoV-2 vaccine.

## Acknowledgments

We gratefully acknowledge the authors, originating and submitting laboratories of the sequences from GISAID’s EpiCov™ Database on which this research is based. We thank Drs. Robert Gramzinski and Lydie Trautmann.

## Author contributions

Conceptualization: M.R., K.M.; Investigation: B.D., E.L., H.B.; Writing – Original Draft: M.R.; Writing – Review & Editing: all authors; Funding Acquisition: K.M.

## Supplementary Data

Supplementary data include Materials and Methods and Figures S1-S5. Sequences and metadata are provided as Supplementary Data file S1.

## Supplementary Data

### Materials and Methods

#### Sequence data

All sequences were downloaded from GenBank or GISAID (https://www.gisaid.org/). GenBank accession numbers are: MN970003, MN970004, MT049951, MT050417, MT111895, MT123292, MT123293, MT127113, MT159778, MT192758, MT226610, MT232871, MT232872, MT256917, MT256918, MT259226, MT259227-MT259231, MT259238, MT259259, MT263459, MT273658, MT281530, MT281577, MT291826-MT291836, MT292580-MT292582, MT293547. Sequences downloaded from GISAID are available in Supplementary Data S1. ‘GISAID_acknowledgement_ table_20200410.xls’.

#### Sequence analysis

All SARS-CoV-2 sequences available as of April 2020 were downloaded and deduplicated. Sequences known to be linked through direct transmission were down sampled and the sample with the earliest date (chosen at random when multiple samples were taken on the same day) was retained. Insertions relative to the reference sequence (Wuhan-Hu1, GISAID accession EPI_ISL_402125) were removed and the 5’ and 3’ ends of sequences (where coverage was low) were excised resulting in an alignment consisting of the 10 open reading frames (ORFs). Sequences were aligned with Mafft v7.453 (*15*). Phylogenies were obtained using FastTree v2.10.1 (compiled with double precision) (*16*) under the GTR model with gamma heterogeneity. Tips more divergent than expected from their sampling date were removed from **Figure 1**. Branches are given in units of substitutions per site and the phylogeny was rooted using the oldest sequence.

Sites under positive, negative, and neutral selection were estimated using a mixed-effect likelihood method that estimates non-synonymous and synonymous substitution rates at each codon (*18*). Non-synonymous and synonymous substitution rates were estimated using a reversible MG94 codon model (*19*). Estimates were drawn from internal nodes only. Genetic differentiation between subpopulations were performed on each gene separately using Nei’s G_ST_ (*21*) and Jost’s D (*22*) with the mmod package (*23*) and calculated over 100 bootstrapped samples.

A neutrally evolving phylogeny was simulated with a constant birth-death rate; a phylogeny with increasing fitness was simulated with exponentially increasing birth rate and constant death-rate. Branch-lengths for both simulated phylogenies were drawn from the SARS-CoV-2 *spike* gene phylogeny. Mutational fitness over time was estimated for each phylogeny by extracting branches from each node independently and calculating the peak height of the spectral density profile of the graph Laplacian of each subtree with >4 tips (*29*). Each phylogeny was down-sampled 100 times at 50% and mutational fitness was re-calculated.

#### Ancestral reconstruction

Ancestral protein sequences were reconstructed on amino acid alignments using maximum posterior probability and returning a unique residue at each site assuming a JTT model with gamma heterogeneity. A sliding window of ten amino acids (and a step of one amino acid) was used to compare the cumulative number of mutations in the Human-bat and Human-bat-pangolin ancestors with respect to the human ancestral sequence. Median values for each window were compared to a null window (computed as a normal distribution of ten values with a mean equal to the mean value across the entire Spike protein, 0.046 mutations) using a one-tailed t-test.

**Figure S1.**
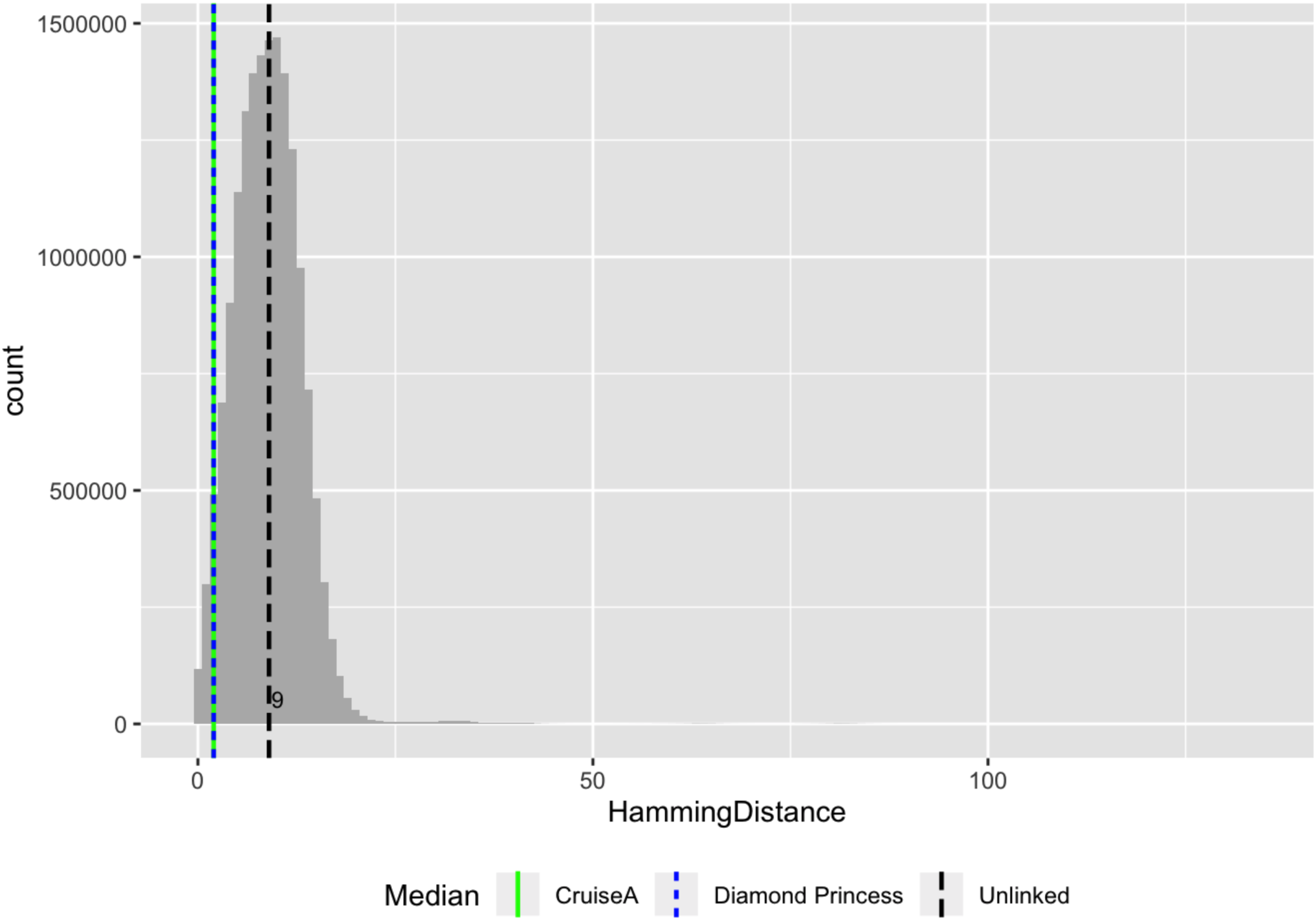
Narrow distribution of hamming distances across SARS-CoV-2 genomes. Hamming distances were calculated across 5,712 genome sequences. The x-axis represents the number of nucleotide mutations between two genomes. The median is reported for all unlinked individuals (n = 5,597, median = 9) as well as for linked cases from Cruise A (n = 25, median = 2) or from the Diamond Princess cruise (n = 69, median = 2).

**Figure S2.**
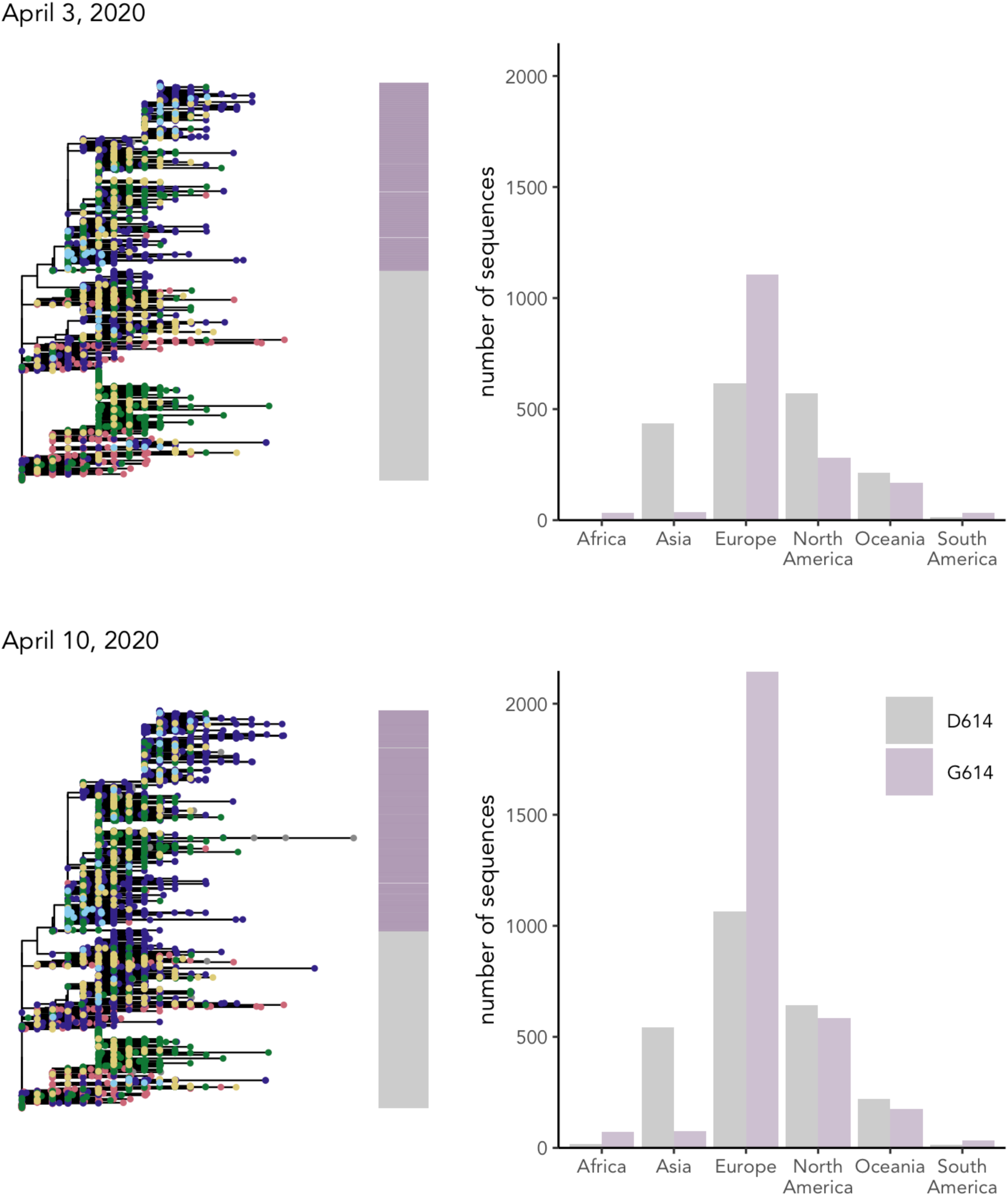
The Spike mutation D614G quickly became dominant. The mutation D614G was found in 55% of sequences sampled globally as of April 10, 2020, the second most frequent mutation in the Spike was only found in ∼1% of sequences. The figure depicts the distribution of D and G variants (upper panel, on April 3, 2020; lower panel, on April 10, 2020). This mutation corresponded to the most frequent variant in Europe, but it was only rarely sampled in Asia. The phylogeny suggests that this mutation was linked to a bottleneck event when SARS-CoV-2 viruses were introduced in Europe; this mutation was first seen in a sequence sampled in Germany at the end of January. There is no evidence that the increasing predominance of this mutation was caused by convergent selection events that would have occurred in multiple individuals.

**Figure S3.**
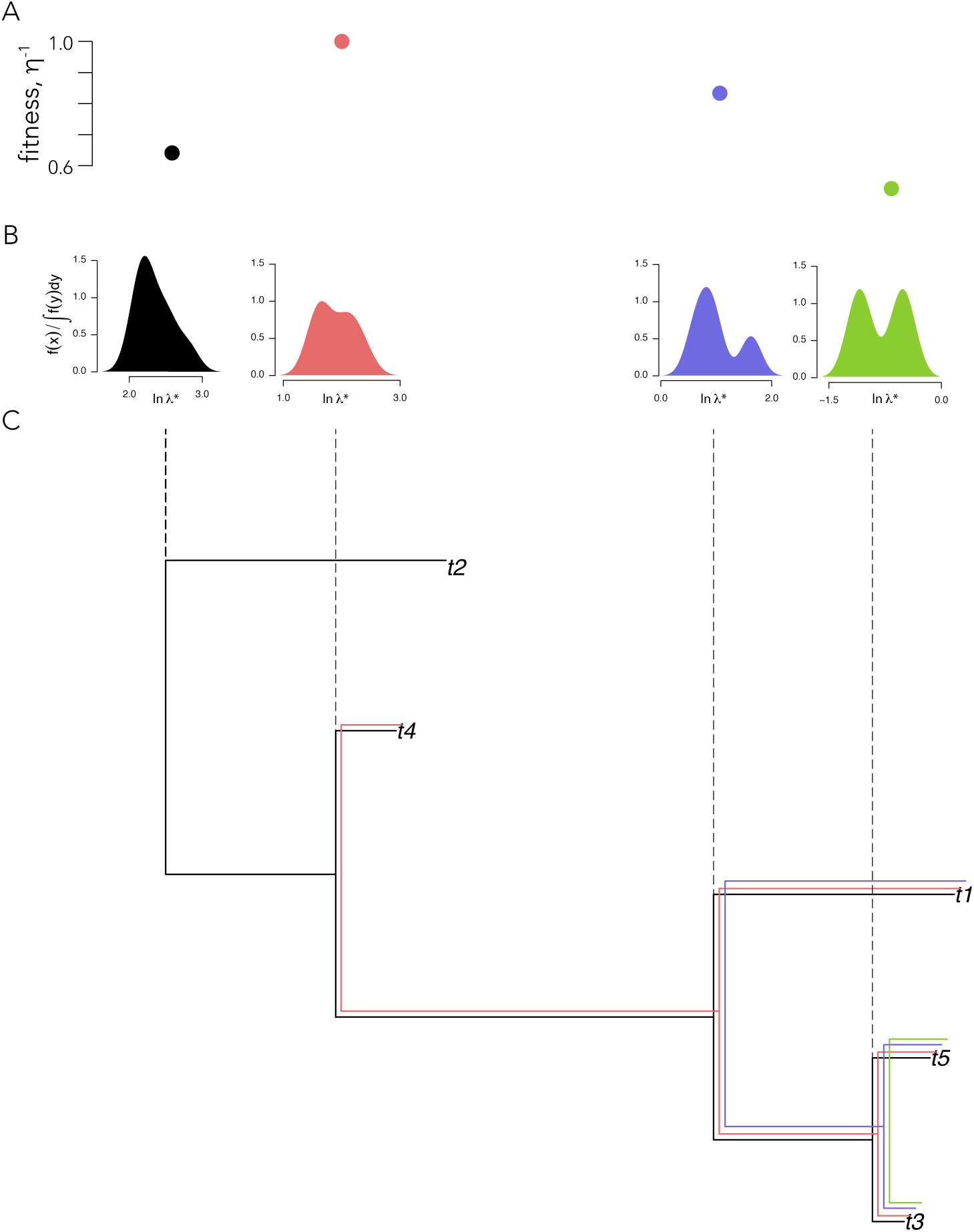
Mutational fitness through time. (A) Mutational fitness was measured as the peak height (1/η) of the (B) spectral density profile of the graph Laplacian constructed from the (C) phylogeny descending from an internal node. The sequence of mutational fitness estimates over time demonstrate the temporal pattern of (un)changing mutational fitness.

**Figure S4.**
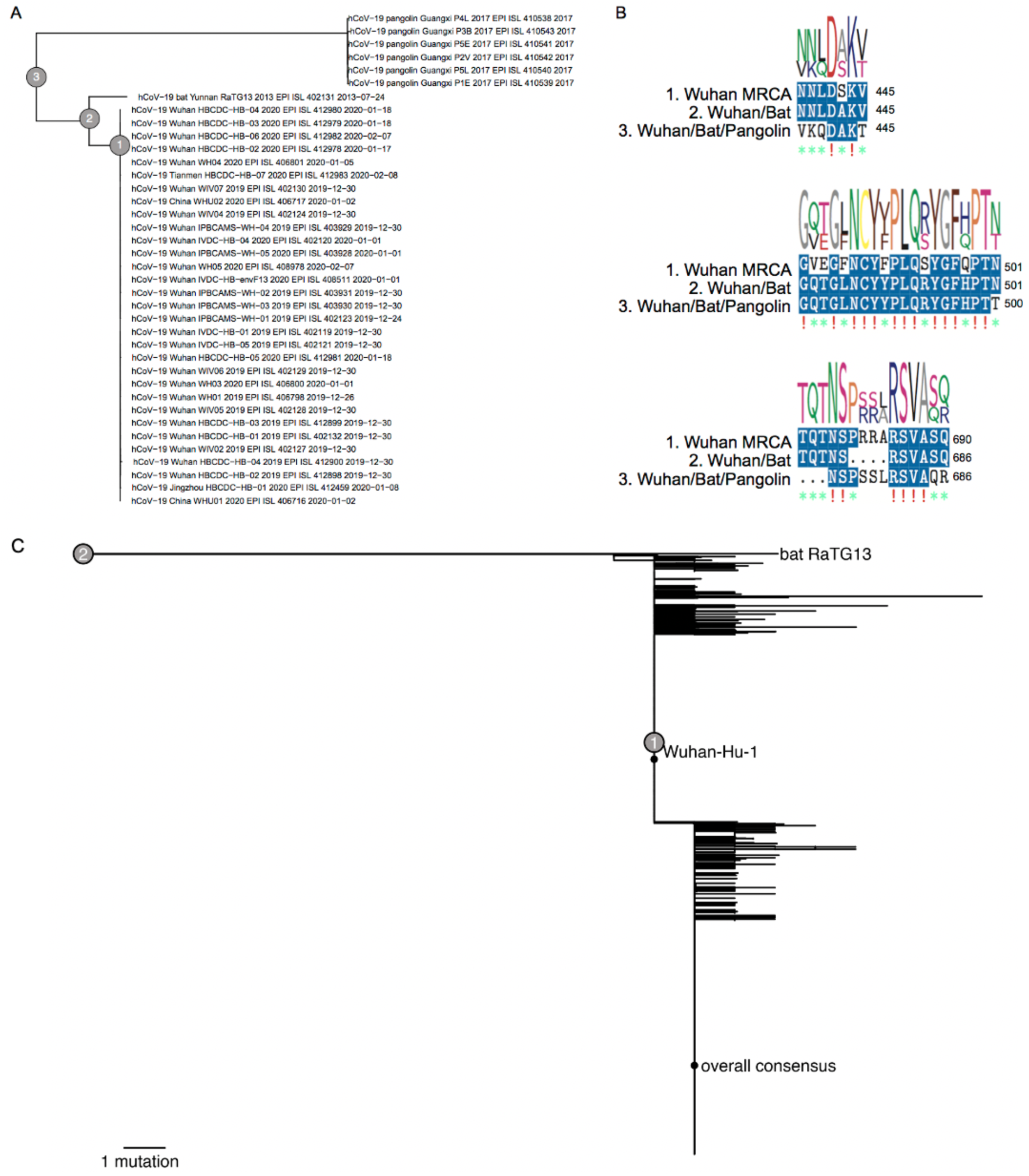
Phylogenetic tree of SARS-CoV-2 spike sequences. (A) Phylogenetic tree reconstructed from SARS-CoV-2 genomes corresponding to sequences sampled from Wuhan, China, seven sequences sampled from Pangolins between 2017 and 2019, and one sequence from a bat collected in 2013. Human (1), human and bat (2), and human, bat, and pangolin (3) ancestral nodes are indicated. (B) Amino acid sequence alignments from the most diverse segments between the three reconstructed ancestral sequences. (C) Phylogenetic tree reconstructed from circulating and optimized vaccine candidate (consensus and MRCA) sequences. The rarity of polymorphisms is evident in the shallowness of the tree. Highlighted are sequences corresponding to the consensus for all circulating sequences and the MRCA for the sequences from Wuhan and for the nodes including sequences from pangolins and bats.

**Figure S5.**
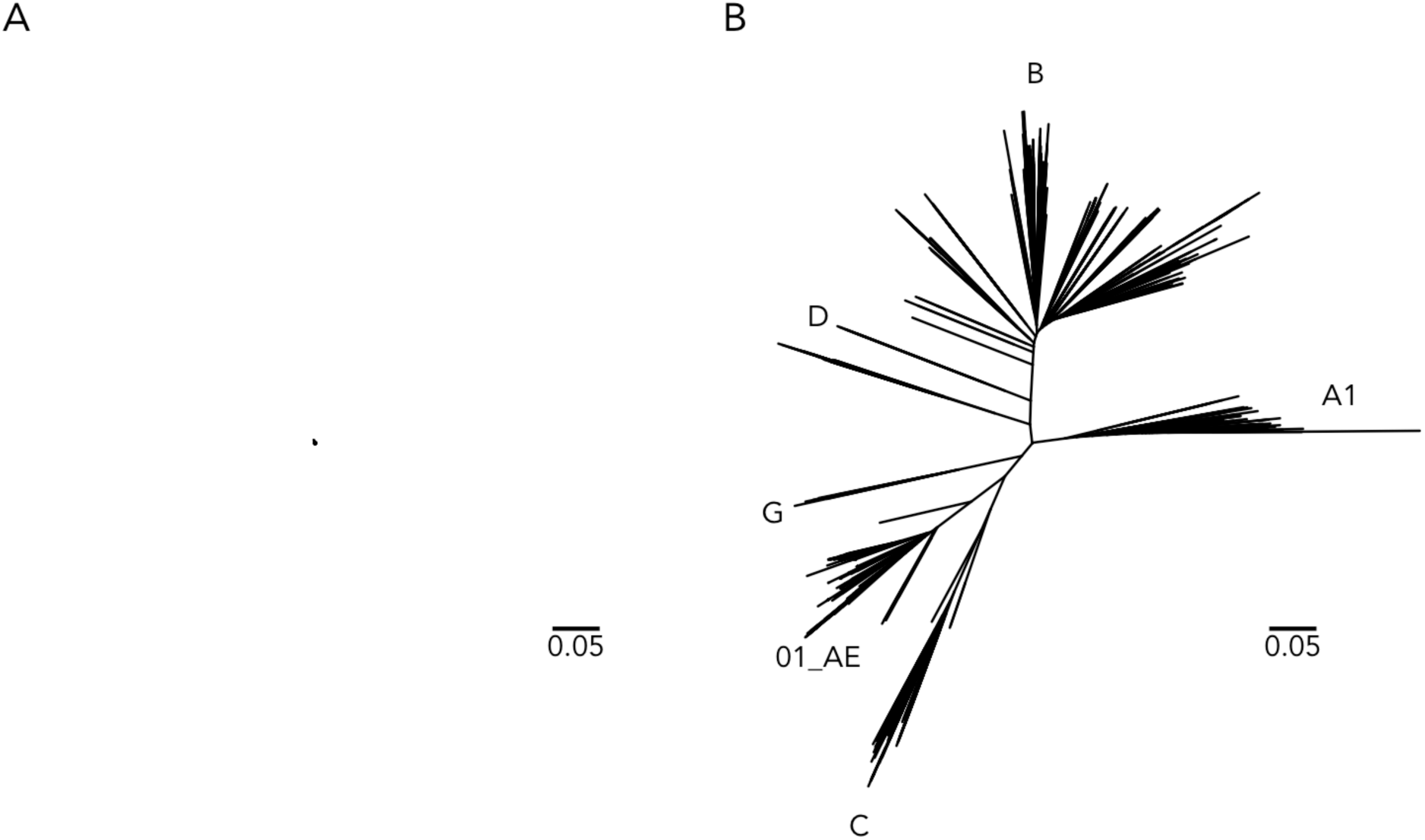
Comparison of the diversity found across circulating SARS-CoV-2 or HIV-1 sequences. The trees show the diversity found across SARS-CoV-2 *spike* sequences (n = 5,579) (A) and across HIV-1 *envelope* sequences sampled in 2010 (n = 1,067) (B). HIV-1 sequences were downloaded from the LANL HIV-1 database (www.hiv.lanl.gov). The purpose of this figure with both trees shown on the same scale is to illustrate the extent of diversity that needs to be covered by an HIV-1 vaccine compared to what a SARS-CoV-2 vaccine shall cover.

